# Quantification of breeding sites parameters in shaping bacterial communities in *Aedes aegypti* (Diptera: Culicidae)

**DOI:** 10.1101/2021.10.24.465617

**Authors:** Josiann Normandeau-Guimond, Lyza Hery, Amandine Guidez, Audrey-Anne Durand, Christelle Delannay, Jean Issaly, Stéphanie Raffestin, Joseph Nigro, Anubis Vega-Rúa, Philippe Constant, Claude Guertin, Isabelle Dusfour

## Abstract

Studies have demonstrated the importance of breeding site, few had disentangled the role of microbiome, physico-chemical and biological factors of water as well as landuse on larval microbial communities and their recruitment in mosquito. A quantitative exploration of the interplay of multiple factors on mosquito microbiome was performed using a dataset obtained through a field survey undertaken in French Guiana. Two complementary hypotheses were tested (i) the most dissimilar larval microbiome structures in breeding sites displayed the most contrasting water properties and land-use, (ii) a higher specificity level of environmental parameters have an incidence on larval microbiome. Variance partitioning approach validated the two hypothesis by providing evidence that water bacterial community is a most significant driver shaping the structure of the bacteriome in mosquito than other environmental parameters from the breeding sites. However, land-use does not play such important role to explain variance. Our results consolidate and complement the knowledge shaping mosquito microbiota but also highlighted the large unknown in understanding the ecology of the recruitment into host.

## Introduction

The study of microbiota in general and in mosquitoes in particular revealed the complexity of adaptation that do not rely on organisms but also on microbial assemblages and their interaction. Microbial ecology has strongly benefited from ‘omics’ advances by the possibility to reveal thorough portraits of all microorganisms in virtually any kind of environment. Microbiome studies have then consequently flourished in several fields of investigation, describing community structures, deciphering associations between microbial communities and many phenotypes, and demonstrating an essential role in organisms life-traits and adaptation. To date, the structure of the intestinal or complete microbiota of mosquitoes has been associated with the stage of development, sex, taxonomic status and genetics, various organs but also with the type of blood meal, blood meal intake, behavior, vector competence or resistance to insecticides ^1–5^. Understanding intrinsic or extrinsic mechanisms of recruitment may become crucial to counteract pathogen transmission, develop novel method of control or set-up more efficient directed manipulations. Mosquitoes’ gut microbiota at larval and adult stages are partially acquired through the water of their breeding site and by the ingestion of microorganisms colonizing the egg shell ^4,6,7^. Community is renewed at each instar and is almost fully eliminated before adult emergence. Additionnally, endosymbionts are inherited from the mother to the egg. Microorganisms thriving in water are recruited by larvae, resulting in more dissimilarity in bacterial community composition in larvae from different collection sites than in larvae collected from the same site^8,9^. Moreover, microbial community structure and evolution are sensitive to biological and physico-chemical properties of the water and the microbial diversity in place ^4,7,10–12^. Landuse and geographiccal positions were then associated to microbiota structure in the environment and in mosquitoes, most likely the consequence of a cascade effect ^7,8,13,14^.

Even if studies demonstrate the importance of breeding site, few had disentangled the role of microbiome, physico-chemical and biological factors of water as well as landuse on larval microbial communities and their recruitment. This article proposes a quantitative exploration of the interplay of multiple factors on mosquito microbiome. A sub-dataset obtained through a field survey undertaken in French Guiana and Guadeloupe, and published in Hery et al.^15^, was used to test two complementary hypotheses. Firstly, we examined whether the most dissimilar larval microbiome structures were found in breeding sites displaying the most contrasting water properties and land-use. Under this assumption, the environment is seen as a strong determinant of larval microbiome. The second hypothesis considers a higher specificity level of environmental incidence on larval microbiome. Specific characteristics of water and land-use features are expected to explain compositional change of larval microbiome amongst the different breeding sites. Under this scenario, microbial communities associated to larvae thriving in breeding sites displaying contrasting characteristics can be similar. Beyond understanding the ecological interaction in the larval – breeding sites ecosystems, such an approach could initiate the identification of key components essential to elaborate or direct mosquito microbiota manipulation and further study specific traits in mosquito biology ^16,17^.

## Material and methods

### Sampling

Collection sites were situated in several localities from urban to more rural settings on the littoral area of French Guiana, South America. We partially used a dataset published in Hery et al. 2020^15^ by focusing on collection sites from French Guiana complemented with 12 more. In consequence, 71 were used in this study out of a total of 138 *Aedes aegypti* breeding sites prospected during the dry season in October and November 2017. Those breeding sites are representative of type diversity across French Guiana^15^. Sites with 3rd to 4th-instar larvae were favoured. For each site, observed environment parameters were recorded (i.e. volume, type of container, material, shadow, presence of nymphs, presence of fragments or living plants) (Table S1, Field observed dataset, upon request) ; some water physico-chemical parameters were measured on-site using a multi-parameter probe (HANNA instruments, France in French Guiana) (i.e. temperature, salinity, conductivity, dissolved oxygen, ORP) (Table S2, Micro dataset, , upon request), and all insects and water (approx. 700 ml) brought back to Institute Pasteur of French Guiana for immediate processing and storage.

### Water sample processing

All physicochemical parameters determined in the laboratory were analyzed according to accredited standard methods (www.cofrac.fr) by the Laboratory of Environmental Hygiene (LHE) at the Institute Pasteur of French Guiana (Table S2, Micro dataset). Dissolved oxygen (mg/L), turbidity (FNU) and chemical oxygen demand (COD) (mg/L) were measured using 200 mL of water immediately when returning to the lab the day of collection. The remaining volume of water sample (~500ml) was frozen at −20°C. At the end of the collection campaign, the samples were sequentially melted, then centrifuged at 8000 rpm, for 10 min, at 4°C. Resulting supernatants were used for determination of a following parameters : Ca+, Mg+, K+, Cu2+, Fe2+, Zn2+ (Table S2, Micro dataset) while the pellets obtained were processed for DNA extraction. Total genomic DNA was extracted from water pellets using the NucleoSpin® Soil kit (Macherey-Nagel, USA) according to the manufacturer’s instructions.

### Larval sample processing

Back to the insectary, larvae were set up in pans. Twenty to 30 larvae of 3rd to 4th instar stages were surface-cleaned according to Zouache et al.^18^ pooled and stored at −20°C. The number of *Aedes aegypti* larvae, other mosquitoes and chironoma were estimated and converted in insect density per liter of water (Table S1 Field observed dataset). In some cases, few days were elapses before harvesting with the objective to let the larvae grow. The pooled larvae were surface-sterilized according to Zouache et al.^18^. Genomic DNA from each sample was extracted according to Hery et al.^15^ using NucleoSpin® Soil kit’s instructions.

### Microbiome data acquisition

The V6-V8 region of the 16S rRNA gene was sequenced using the primers B969F-CS1 5’-ACGCGHNRAACCTTACC-3’ and BA1406R-CS2 5’-ACGGGCRGTGWGTRCAA-3’^19^. The sequencing technology used was Illumina MiSeq 250 bp paired-ends at the Quebec Genome Innovation Centre (McGill University, Montreal, QC, Canada). Sequences were processed using the software USEARCH following the UPARSE pipeline^20^. The paired-end sequences were merged with a minimum length overlap of 400 bp and a maximum overlap of 500 pb. The maximum allowed ratio between the number of mismatched base pairs and the overlap length was set to 0.3. Reads with low-quality scores were removed using a maximum expected error value of 1.0. The remaining high-quality reads were de-replicated, sorted by size, and all singleton sequences were removed. The remaining reads were clustered into operational taxonomic units (OTU) with the UPARSE OTU clustering method using a 97% identity threshold. Taxonomic assignment was realized with the Ribosomal Database Project (RDP) classifier version 16 to remove OTU identified as archaea and chloroplasts. Sequences represented by less than 0.005% of the total number of reads per library were removed. The final taxonomic assignment was done with RDP with a minimum of 80% confidence level.

### Landuse and cadastre datasets

The larger environmental dataset was extracted from available data collected via satellite images analysis and freely available.This includes road network (source: openstreet map contributors), hydrographic network (source : IGN 2019) and land use (source : ONF-2015) and cadaster (available at cadastre.gouv.fr). We used ArcGIS software to extract 27 parameters of quantitatives variables in 50m and 100 m radius : surface of landuse and cadaster, length of roads and rivers, and distance to class of forest, to seashore, to road or to hydrographic network (Table S3, Macro dataset, upon request).

## Data analysis

### Data transformation and matrices set-up

Seventy-one sites presenting all the variables were kept to produce 5 data matrices and GPS positioning files : OTU in water (Table S4, , upon request), OTU in larvae (Table S5, , upon request) and the three environmental dataset listed above. Sparse, zero-dominated compositional datasets (i.e. microbiome matrices and landuse matrices) were box-chord transformed^21^. Other quantitative data were standardized. Variables having a collinearity above th=0.9 were removed using the vif function from the ‘vegan’ package^22^ in the R environment^23^.

## Variation Partitioning

To evaluate how much the structure of larval communities as a whole are explained by water microbiome and environmental variables, we proceeded to variation partitioning analyse^24^ using two independant paths. The first approach was computed to test the hypothesis that the most dissimilar microbiome structures were observed in the most contrasting breeding sites. Identification of the most important gradients distinguishing larval and water microbiome structures, water biophysicochemical properties and land-use were defined by unconstrained Principal Component Analysis (PCA)^25^ or Factor Analysis for Mixed Data (FAMD)^26^, according to variable type using the factoextra R package^27^. The five first components of the ordination analyses were considered as the most important gradients distinguishing breeding site and larvae characteristics. The relationship between larval microbiome gradients and those observed for environmental variables was then analyzed through RDA. Parcimonious models were obtained by using the OrdiStep function (forward direction) from the ‘vegan’ package. The most significant variables of individual RDA were included in the variation partitioning approach computed with the varpart function in the ‘vegan’ package. The second approach tested the hypothesis that the relationship between larval microbiome and breeding sites environmental features is specific to certain factors that are not necessarily the most discriminant amongst sampled sites. Constrained redundancy analyses (RDA) were performed to explain variations of larval microbiome with water microbiome, water physicochemical properties and land-use.

Parsimonious models explaining the best the data were included in the variance partitioning approach.

## Results

### Data overview

Among the 71 sites included in the analysis, 906 bacterial OTU were retrieved from which 817 were found in larvae and 878 in water. In details, 789 were common to water and larvae, 29 were only found in larvae and 89 were only found in water. A total of 121 variables from the 878 input variables in water and 157 variables from the 817 in larvae had collinearity problem. In the water chemistry dataset, salinity out of 16 variables was removed; 9 parameters out of 27 were in the landuse matrix and none out of 19 in the field observed dataset.

### Extreme environmental values are coupled to larvae sites displaying the most dissimilar bacterial composition

The five first axis of non-constrained analysis of individual matrices represent 25.54%, 25.21%, 66.37%, 48.43%, 66.17% of cumulated variance for respectively larval OTUs, water OTUs, water physico-chemistry, field observed data and landuse matrices. After ordistep forward procedure, individual RDA revealed first that the dimension 2, 3 and 4 from PCA of water OTUs provide the better model to explain 17.76% the variance of those from larval OTUs (Table 1). Second, only one dimension of PCA water physico-chemistry parameter provides the best linear model and explains 7.51% of variance in larval OTUs (Table 1). Thirdly, dimension 1, 3 and 4 from observed data FAMD explained 7.49% of larval variance extreme values (Table 1). Finally, no significant model were obtained from landuse-based analysis (Table 1).

**Table 1 :**
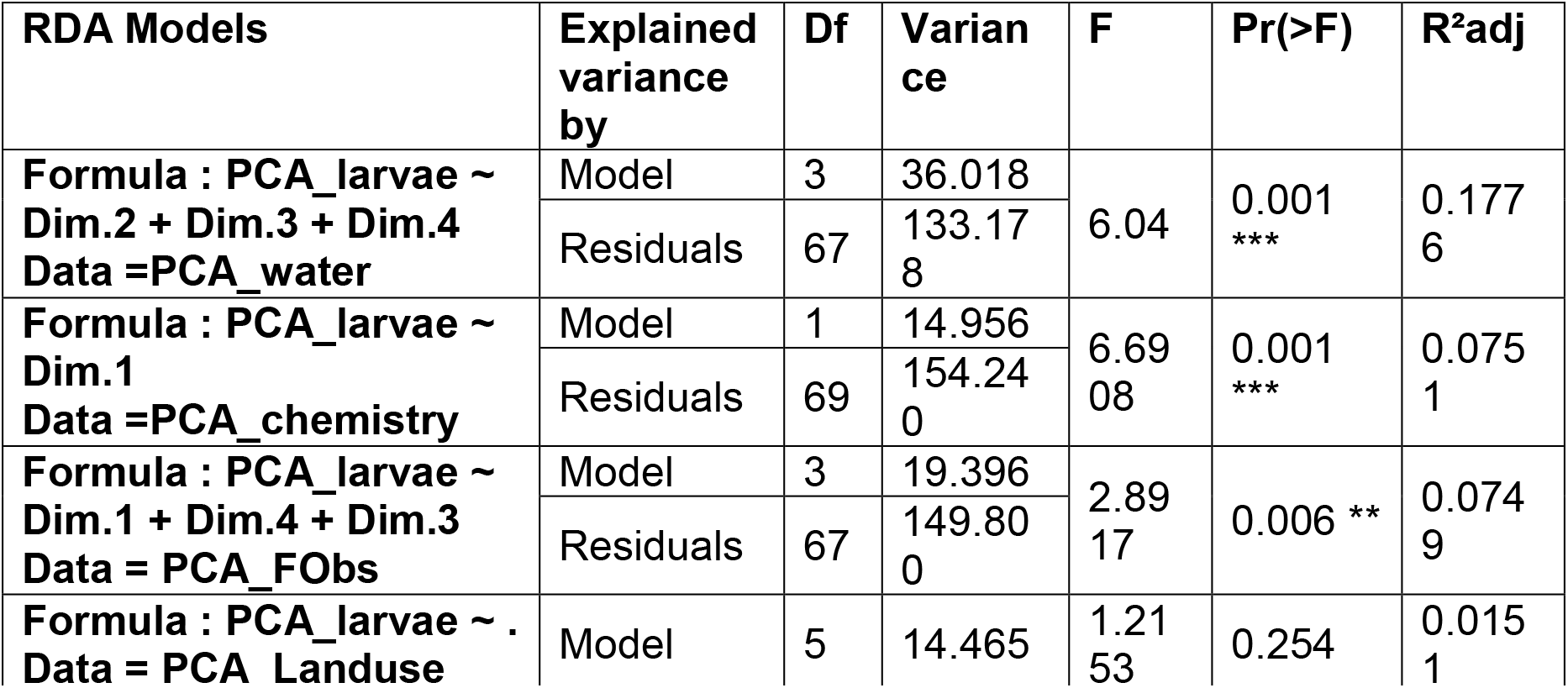
results of individual rda models for each explanatory variables coordinates matrices against larval one.

| **RDA Models** | **Explained variance by** | **Df** | **Variance** | **F** | **Pr(>F)** | **R<sup>2</sup>adj** |
| --- | --- | --- | --- | --- | --- | --- |
| **Formula : PCA_larvae ~ Dim.2 + Dim.3 + Dim.4**<br>**Data =PCA_water** | Model | 3 | 36.018 | 6.04 | 0.001<br>*** | 0.177<br>6 |
|  | Residuals | 67 | 133.178 |  |  |  |
| **Formula : PCA_larvae ~ Dim.1**<br>**Data =PCA_chemistry** | Model | 1 | 14.956 | 6.6908 | 0.001<br>*** | 0.075<br>1 |
|  | Residuals | 69 | 154.240 |  |  |  |
| **Formula : PCA_larvae ~ Dim.1 + Dim.4 + Dim.3**<br>**Data = PCA_FObs** | Model | 3 | 19.396 | 2.8917 | 0.006 ** | 0.074<br>9 |
|  | Residuals | 67 | 149.800 |  |  |  |
| **Formula : PCA_larvae ~ .**<br>**Data = PCA_Landuse** | Model | 5 | 14.465 | 1.2153 | 0.254 | 0.015<br>1 |

Overall extreme environmental values explained almost 23% of extreme dissimilarity in bacterial communities from larvae. Variation partitioning approach demonstrated that the participation of the breeding site bacterial community alone reaches 10.57% in our dataset, largely above the rest of included parameters (Figure 1). Also, the variation explained by two or three matrices together is more or as much as important as water physico-chemistry or field observed ones (Figure 1).

**Figure 1 :**
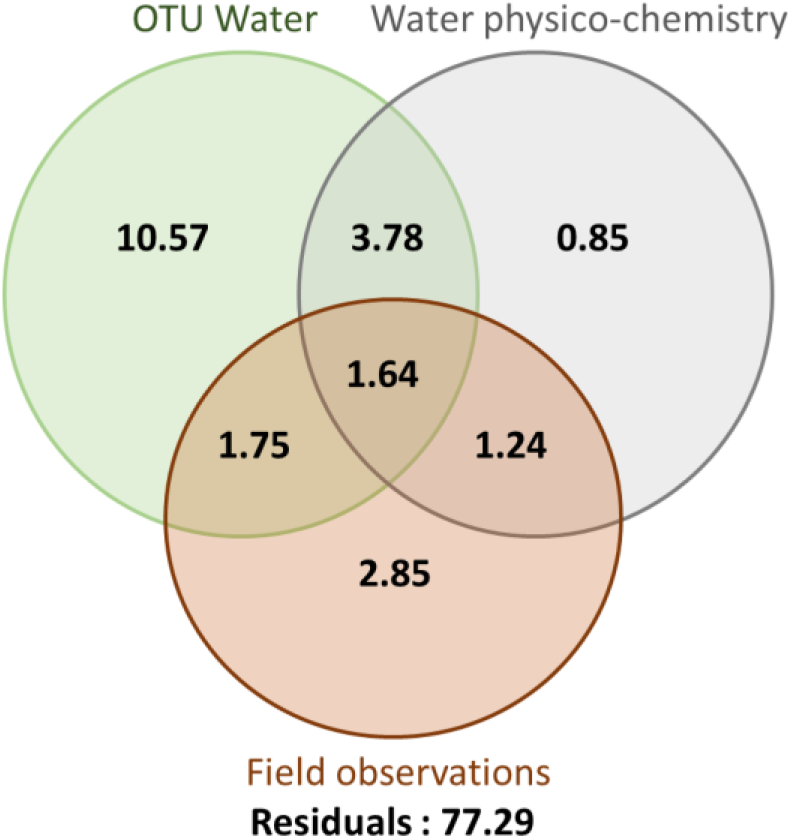
percent explaining the variance in the larval bacterial community for each and associated environmental matrices obtained through variation partitioning analyse

The most divergent gradients of environmental features distinguishing the breeding sites were identified though unconstrained principal component analyses. We quantified for the first time that extreme environmental values are coupled to breeding sites harboring larvae displaying the most dissimilar composition. This field survey quantitatively confirms that larvae thriving in ecologically contrasting breeding sites are expected to harbor different microbiomes.

### Small set of environmental variables explained 18% of variance in larval bacterial communities

The second approach aimed at determining which environmental factors impact larval bacterial composition. To do so, individual RDA constraining larval OTUs with each environmental matrix were performed to model the dataset. A primary RDA was performed with OTU characterized in water. Based on that analysis, OTUs within the 1% highest and lowest biplot scores were extracted counting for 36 OTUs. The subsequent RDA explained 13.56% of variance. After the ordistep selection, a model composed by eleven OTUs and explaining 11.18% of variation was obtained (*p-val*= 0.001, R^2^Adj = 0.1118). Among them OTUs from the Alphaproteobacteria class were majoritary (N=5), followed by Actinobacteria one (N=3), Cyanobacteria (N=1) and Sphingobacteriia (N=1). One OTU was not identified at the Class taxonomical level. RDA computed with physico-chemical data revealed that a model composed by Turbidity, ORP (Oxido-reduction potential) and Calcium ion (*p-values* = 0.001) explained 2.67% of variance. In addition, only the volume of containers explain 1.35% of variance when constraining the larval dataset with the field observed variables. Finally, the surface of sea within a 50m-radius, distance to the road, surface of forest in a 100m-radius significantly explain 3.95% of the variance in the final model (*p-value* = 0.001). Computing the variance partitioning analysis showed that 17.59% of variation is explained by the four matrices (Figure 2). Eleven OTUs in water explained 10.87% by itself ; Ca+ concentration, turbidity and ORP explained 2.51% ; volume explained 0.17% and surface of sea ; distance to road and surface of forest explained 1.64%. The remaining 2.4% are composed by interaction between datasets.

**Figure 2 :**
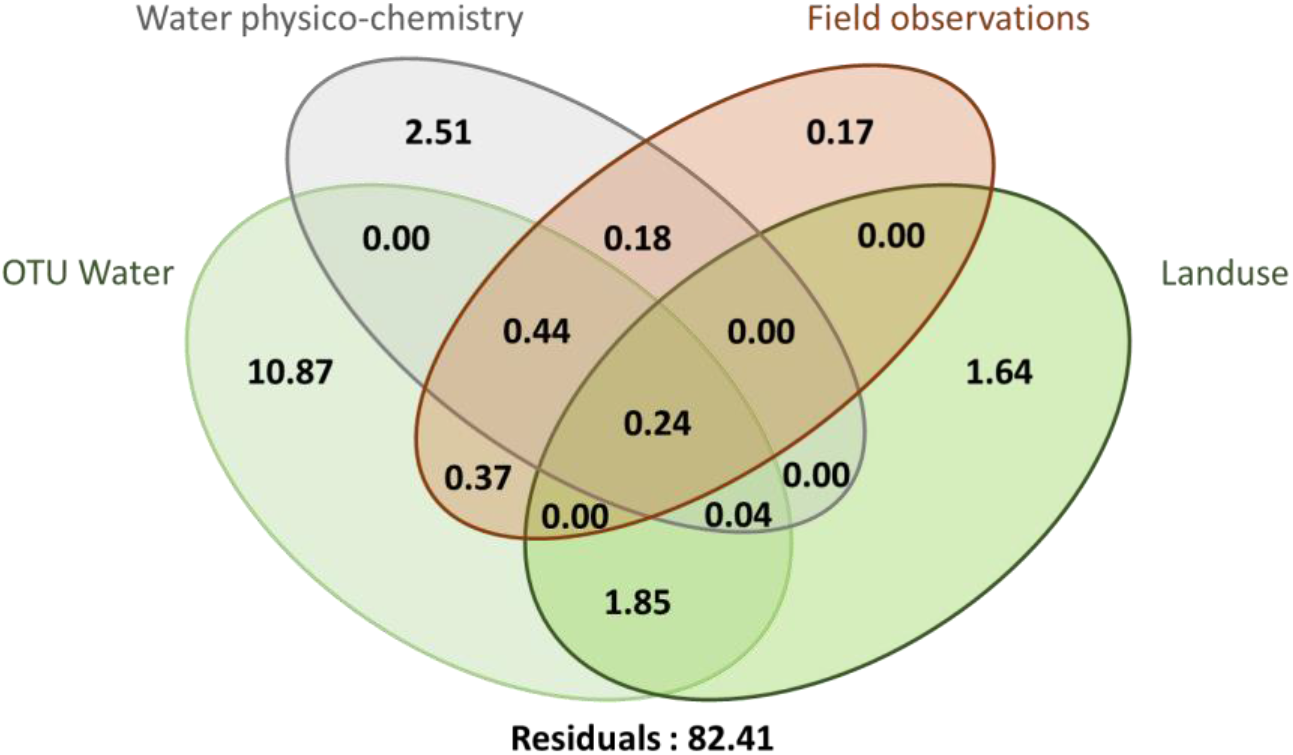
percent explaining the variance in the larval bacterial community for each and associated environmental matrices obtained through variation partitioning analyse

Those results are congruent with the non-constraint approach showing that bacterial microbiome play the main role amongst all the environmental datasets tested but also highlight that above 77.29% of variance remains unexplained by the paramaters we included (Figure 3).

**Figure 3 :**
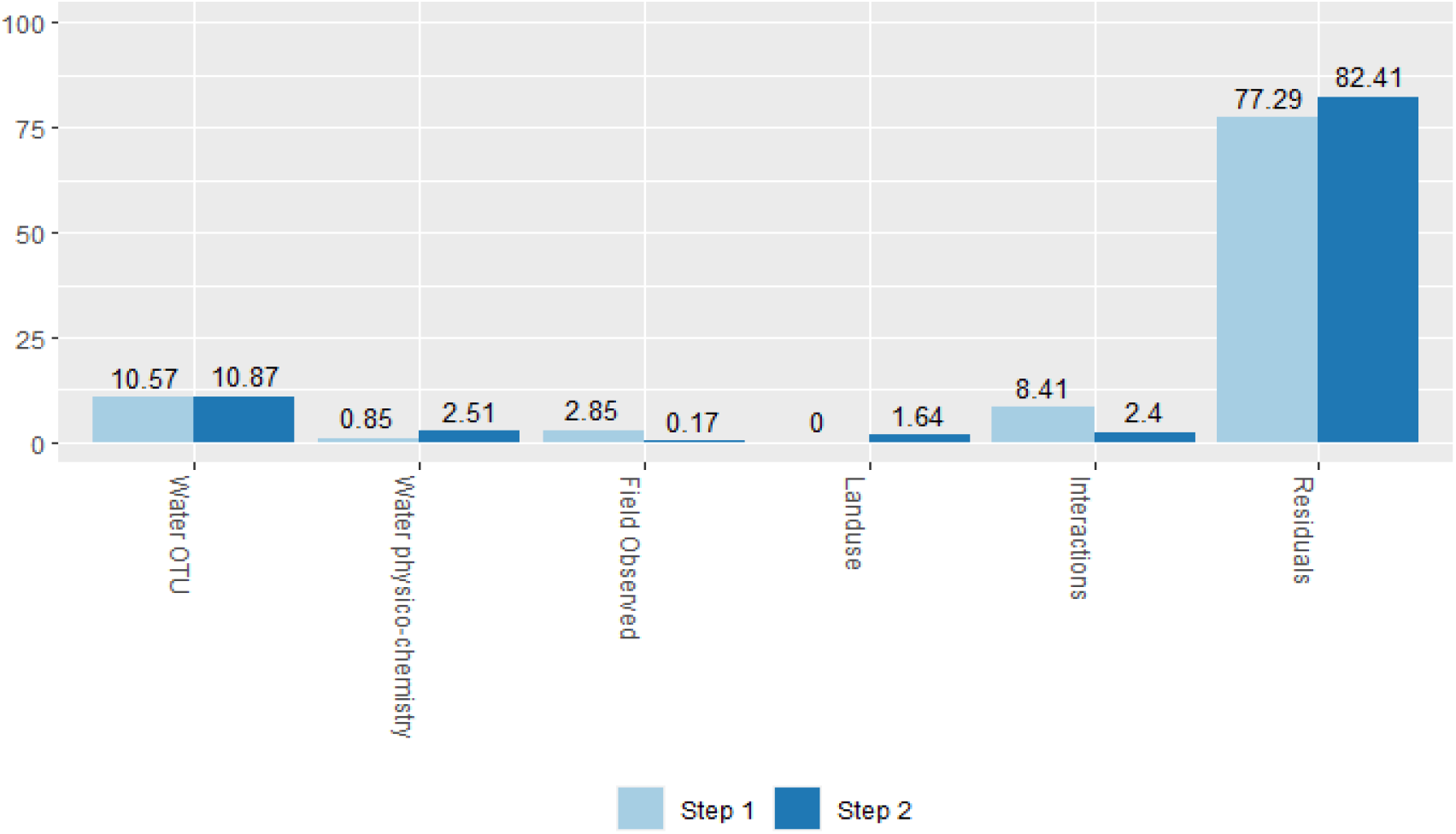
barplot of percent of percentage of each matrices explaining larval data set in step 1 and 2 of the analysis.

## Discussion

The main objective of our study was to disentangle the contribution of environmental parameters in explaining variation of mosquito microbiome structure among different BS. Two complemetary hypotheses were tested using either unconstrained or constrained multivariate analysis combined with variation partitioning^25^. The first hypothesis assumed that the most dissimilar larval microbiome structures are found in breeding sites displaying the most contrasting water properties. We extracted the most extreme components of our datasets through non-constrained multivariate analysis and then validated our assumption. Beyond the close associations between microbiota and their environment that were previously observed^4,6,7 10^, we demonstrated that only closely related parameters such as water microbiome assemblages, water physico-chemical parameters and observed data in the surroundings exert a structuring effects and explain 22.69% of variance in mosquito microbiota. Land-use in a radius up to 100m did not participate to extreme dissimilarity in larval assemblages. Amongst datasets with the strongest explicative power, water OTU is remarkable by explaining 10.57% of mosquito bacterial OTU variance (46.58% of explained variance), followed by field observation that explained 2.82% (12.42%) and water physico-chemistry corresponding to a marginal 0.85% (3.95%). Even though water physico-chemistry was not directly associated with extreme patterns of mosquito microbial structure, variation partionning highlighted redundancy amongst datasets.The analysis revealed that this role is mainly in interaction with water bacterial communities (3.78%) and with field observations (1.24%). This may be explained by the direct role of water components in structuring aquatic microbe assemblies then eventually on the recruitment into the mosquito. It is indeed acknowledge that such a structure is related to its environment and impacts all aquatic organism microbial communities^28,29^. Field observations composed a set of heterogeneous biological and physical visible parameters that were recorded at the breeding sites.Various interactions could occur with this dataset as it covers volumes, colors, shade, material types, presence of vegetation, number of other insects etc. Interactions are always difficult to interprete and must be taken carefully.

The second hypothesis was tested by modelling the bacterial communities structures with variables from each datasets matrices. Those specific characteristics explained up to 19% of variance, in accordance with the first approach. All datasets explained a part of the variance in the mosquito dataset, including land-use characteristics suggesting that they could modulate the structure at finer scale. Eleven bacterial OTUs from water structured and explained alone 10.87% (46.54% of the explained variance) variation of bacterial OTUs distribution pattern in larvea. Taxonomic identifications possible at the Class level are in congruence with other publications on mosquito or water microbiome studies^15,30^. Turbidity, Oxido-reduction potential and Calcium ion were the variable contributing the most in explaining variation of larval microbiome structure (11.43% of the explained variance). All those parameters are known drivers of microbial assemblages. Related to the presence of particles in a fluid making water opaque^30,31^, turbidity reduces the availability of photosynthetic active radiations for photosynthetic bacteria and algea, alterating the trophic network of microorganisms. Oxido-reduction potential is altered by pH, known as one of the most important feature shaping microbial communities. Finally, ion calcium is a key component for bacteria functions, including biofilm formation^32,33^. Within the field observed data, only volume of BS explained 0.17% (0.72% of the explained variance) of variation of density of mosquitoes or other insects. This value is marginal and no explanation can yet be assumed. Land-use had been associated to microbial structure in soils but can underly a large panel of factors. The components we extracted (i.e. surface of sea within a 50m-radius, distance to the road, surface of forest in a 100m-radius) explained 1.64% (7.02% of the explained variance). The remaining 8.4% (35.96%) are composed by redundancy between datasets. The panel of factors that are influencing bacterial communities in mosquito larvea is in congruence with other studies and the biology of bacteriome. Both statiscal approaches, targetting either extreme gradients or most influencial variations, converged to the same conclusion with 20% of variation explained by environmental features of BS. These finding supports the notion that BS encompassing contrasting water properties are expected to display dissimilar microbial communities, with a direct incidence on microbiome structure of larvea. Eventhough a long way still exist toward efficiently manipulating environmental parameters of BS to promote the assembly of beneficial microbiota and associated functions, manipulation of turbidity, redox potential and calcium is a promising approach.

Despite intense efforts in characterizing water and landscape attributes of BS, between 77 and 80% variation of larvae microbiome remained unexplained. Inclusion of more parameters should be considered. For instance, mother-progeny inheritance or eggshell microbiome ingestion were not explored ^4,6,7^ ; all other microorganisms living in water and in mosquitoes were not taken into consideration ^8,34^; the temporal trends of colonisation, meteorological data and dynamics of the breeding sites should be worth considering ^29^. In addition, eventhough genetics did not seem to affect bacteria recruitment in absence of selective pressure^7,13^, no current research have demonstrated if it could play a role in recruitment under selective conditions (ex: xenobiotic pressure, food depletion,…).

## Conclusion

This study provides evidence that water bacterial community is a most significant driver shaping the structure of the bacteriome in mosquito than other environmental parameters from the breeding sites. However, land-use does not play such important role to explain variance. Our results consolidate and complement the knowledge on mosquito microbiota but also highlighted the large unknown in understanding the ecology of the recruitment into host.

## Acknowlegment

Authors want to thank the inhabitants of French Guiana for allowing us to sample around their homes.

## Access and Benefit-sharing

In accordance with Article 17, paragraph 2, of the Nagoya Protocol on Access and Benefit-sharing, an internationally recognized certificate of compliance with the number ABSCH-IRCC-FR-246988-1 was issued by the French Government. It permits the utilization of genetic ressources from *Aedes aegypti* collected in French Guiana.

## Funding

This work was supported by Action Concertées Inter Pasteuriennes (Grant ACIP 01-2016) and “an Investissement d’Avenir grant of the Agence Nationale de la Recherche” (CEBA: ANR-10-LABX-25-01).

## Notes

### Competing Interest Statement

The authors have declared no competing interest.

